# Optimizing core collections for genetic studies: a worldwide flax germplasm case study

**DOI:** 10.1101/2025.06.30.662343

**Authors:** Matthieu Gouy, Matthieu Bogard, Faharidine Mohamadi, Boris Demenou

**Affiliations:** ARVALIS Institut du Végétal, 110 chemin de la côte vieille, 31450 Baziège, France; ARVALIS Institut du Végétal, Station Expérimentale, Route de Malesherbes, 91720 Boigneville, France; ARVALIS Institut du Végétal, 45 voie Romaine - Ouzouer-le-Marché, 41240 Beauce La Romaine, France

**Keywords:** Core-collection, optimization criteria, Quantitative Trait Loci (QTLs), genetic diversity, flax (*Linum usitatissimum* L.)

## Abstract

Core collections provide a strategic approach to reducing population size while retaining genetic diversity and allele frequencies, serving as key resources for genetic research. Although various sampling and selection strategies have been proposed, most of them focused on either diversity or representativeness, rarely both, and none fully integrated these with QTL detection optimization. The first part of our study focuses on a genetic diversity analysis of a flax germplasm (*Linum usitatissimum* L.), a prerequisite for the development of core collections. This germplasm, maintained by the Arvalis Institute, is a worldwide flax collection comprising 1,593 accessions originating from 42 countries, encompassing all major flax-growing regions. It includes both spring- and winter-type lines, as well as oilseed and fiber types. The results revealed a pronounced genetic structure within the germplasm, strongly influenced by cultivation purposes (fiber *vs.* oilseed flax), growth cycle (winter *vs.* spring), and geographic origin. A K-means clustering procedure identified six clusters as the optimal structuration, which aligns with our knowledge of this germplasm. Overall genetic diversity was moderate (Hₑ = 0.22), with oilseed flax clusters displaying greater diversity than fiber flax, likely due to the broader selection history and wider geographic distribution, findings consistent with previous studies. In a second step we evaluated twenty distinct strategies for core-collection development. Some approaches were originally developed for core-collection construction while others were developed for optimizing genomic-selection calibration panels. QTL detection performance was assessed via extensive simulations of QTLs distributed across the genome. We observed a fundamental trade-off between maximizing diversity and ensuring representativeness in core collection design. Diversity-oriented approaches may overemphasize rare or outlier genotypes, compromising representativeness, while representativeness-focused strategies leaded to overlooking rare alleles, thus limiting diversity. In our results we have found that particular combinations of selection criteria achieved a favorable balance between genetic diversity and representativeness, while concurrently maintaining a robust capacity to capture QTL signals across the genome. We demonstrated that using the Shannon index combined with the allelic coverage led to optimal core-collection design adapted for GWAS applications in a structured population. These results provide knowledge for the development of optimized core collections tailored to GWAS applications.

## 1 Introduction

Plant genetic resources are a crucial source of diversity and are essential for improving crops. Useful plant genetic ressources for breeding includes landraces, breeding lines, cultivars and the wild relatives of the target species, offering a broad range of alleles that can be exploited to enhance key agronomic traits. To face climate change, food security challenges, and the need for sustainable agriculture, the management and conservation of genetic resources are fundamental for the development of productive and resilient genetic material (Esquinas-Alcázar, 2005). More than seven million plant accessions are conserved across around 1,750 genebanks, with over 60% belonging to only thirty cultivated species (Hammer et al., 2003; Thormann et al., 2012). The diversity maintained is largely oriented towards human needs. Given the high number of accessions to be conserved, it has become necessary to rationalize the management of these resources, particularly through the selection of core collections.

The core collection concept was formalized in the 1980s to ensure optimal management and use of the genetic resources collected over time. A core collection can be defined as a reduced set of accessions that represents the genetic diversity of a species and its wild relatives with minimal redundancy (Brown and Clegg, 1983; Frankel, 1984). Since its inception, numerous studies have focused on methodologies for creating core collections. Brown and Frankel (1989) suggested that a core collection should not exceed 10% of the full collection and should never include more than 2,000 entries. In practice, most core collections represent between 5% and 20% of the original germplasm (Van Hintum et al., 2000). The reduced size of a core collection is crucial to ensure its efficient long-term management. The creation of core collections addresses two main objectives: (1) maximizing the genetic diversity, often favored by taxonomists, geneticists, and gene bank curators, and (2) maximizing the representativeness of the germplasm, typically chosen by breeders (Marita et al., 2000). The goal for the former is to maximize diversity criteria and conserve the rarest alleles. The second objective involves faithfully representing the source germplasm by retaining more generalist alleles. Jointly optimizing these two objectives ensures efficient short- and medium-term management of a species’ genetic resources, although this remains challenging.

Initially, passport information (i.e., morpho-descriptives data, geographical origins) and other phenotypic traits (e.g. earliness, disease resistance traits) were used to establish core collections. However, it was recognized that environmental factors could influence these variables, leading to inaccurate representations of heritable genetic diversity (Tanksley & McCouch, 1997). Nowadays, the use of molecular markers, such as RAPDs (Marita et al., 2000), SSRs (Soto-Cerda et al., 2013), or SNPs (Bianchi et al., 2020; Fu et al., 2025) has become standard and essential for studying genetic diversity and developing core collections.

Many approaches to developing core collection (CC) have been described. For these approaches, a comprehensive characterization of the species’ genetic diversity and structure is an essential prerequisite, as it is critical to ensure that all genetic clusters are adequately represented within the selected subsets of individuals. This requirement underpins the rationale for employing stratified sampling methods, which offer a more suitable alternative to random sampling by preserving the underlying genetic structure (Charmet and Balfourier, 1995; Gouesnard et al., 2001; Franco et al., 2003). The sampling rate (i.e., allocation) must be defined based on the intended objectives. Several strategies have been suggested: a fixed selection rate, independent from cluster size, a rate proportional to cluster size, a logarithmic-proportional rate (which helps maintaining a manageable collection size), or a rate proportional to intra-cluster genetic distances (or other diversity metrics), also known as the D-method. This latter method has shown significant efficiency compared to alternative approaches (Franco et al., 2006).

The selection of accessions can be based on one or several objective(s) (rarely greater than two) to be optimized, such as genetic distances (Jansen and Van Hintum, 2007), diversity criteria (Franco et al., 2006; Thachuk et al., 2009), or the effective alleles number and their coverage rate (Kim et al., 2023). Some strategies have been developed to simultaneously optimize multiple criteria (Odong et al., 2013; De Beukelaer and Davenport, 2018). These approaches rely on optimization algorithms (e.g., genetic algorithms, simulated annealing) which iteratively optimize an objective function (maximizing or minimizing) by picking a new entry, often randomly, at each iteration.

Similar methodologies have been developed in the genomic selection area. These involve the use of calibration set optimization algorithms, which aim to maximize genomic prediction accuracy based on molecular marker data (Laloë, 1993; Albrecht et al. 2011; Pszczola et al. 2012; Rincent et al., 2012; Akdemir et al. 2017; Ou and Liao 2019). While this approach does not directly link genomic selection calibration methods to core collection inception, the optimization techniques used in genomic selection, such as genetic algorithms and diversity-based criteria, could potentially be adapted for core collection creation. The focus on optimizing subsets for prediction accuracy in genomic selection parallels the goal of selecting representative subsets in core collection creation, suggesting a possible methodological crossover. Moreover, core collections are widely used in associations studies for QTL discovery (Nicolas et al. 2016, Berkner et al. 2024). This type of population typically harbors greater genetic diversity than biparental populations and includes a higher number of recombination events. As a result, the resolution of detected QTLs is significantly improved (Breseghello and Sorrells, 2006; Zhao et al., 2007; Huang and Han, 2014; Bandillo et al., 2015).

The quality assessment of a core collection should, whenever possible, be based on data that were not used for its development (Van Hintum et al., 2000). Core collections are often compared to the whole collection (WC) from which they were derived. Various evaluation criteria can be computed to assess the resulting population such as genetic distances, diversity indices (Shannon index, heterozygosity) or even Principal components analysis (Mohammadi and Prasanna, 2003, Reif et al., 2005, and Odong et al., 2013).

The first breeding and improvement flax (*Linum usitatissimum* L.) programs were initiated in the 1920s by Irish and Dutch researchers (Doré, 2006). Breeding efforts specifically targeting fiber flax also began during this period, with early hybridizations carried out in 1919 (e.g., with the EGBK or CRGH lines) (Blaringhem, 1926). Genetic improvement priorities in flax vary according to its intended use, fiber or oilseed, and are primarily aimed at addressing current agronomic and climatic constraints. In fiber flax, breeding efforts focus on enhancing resistance to major fungal pathogens, including *Polyspora lini*, *Septoria linicola*, fusarium wilt, flax scorch, and powdery mildew. Improving resistance to lodging is also a key objective, as it contributes to reducing yield losses and facilitating mechanized harvesting. Additionally, the enhancement of fiber quality remains a central goal, along with the development of cultivars with improved tolerance to abiotic stresses such as elevated temperatures, drought, but also cold, particularly for winter-type lines. For oilseed flax, breeding efforts are focused on stabilizing and optimizing yield while accounting for strong genotype-by-environment (G×E) interactions. Disease resistance, particularly against septoria, is another major goal. Lastly, improving oil quality and enhancing cold tolerance for winter-type lines are key breeding targets. The use of extended genetic diversity in breeding programs could help improving flax for resistance/tolerance to biotic and abiotic factors.

The worldwide diversity of cultivated flax and its wild relatives is represented by an estimated 48,000 accessions maintained in 33 genebanks, among which only around 10,000 are considered genetically distinct or truly unique (Diederichsen, 2007). From these resources, many flax core-collection have been created (Fu, 2006; Diederichsen et al. 2013; Hoque et al., 2020) in order to investigate for example flowering time (Chandrawati et al., 2017), agronomic, seed and fiber quality, disease resistance traits (You et al., 2017), or even powdery mildew resistance (Speck et al., 2022). The Arvalis Institute, a French institute for applied research in agriculture, maintains a collection of around 1,650 fiber and oilseed flax accessions. This germplasm comprises accessions from countries worldwide where fiber and oilseed flax have been cultivated or are naturally distributed, with a particular focus on recently improved lines from western Europe breeding programs. However, no core collection based on this western European flax genetic resources was available. Then, rare genetic studies in Western Europe have examined a diversity panel including large modern Western European flax. In particular, Speck et al. (2022) used a flax panel of 311 lines selected from 38 countries spanning all continents and diverse worldwide climatic regions. However, they did not describe a clear selection methodology to ensure that genetic diversity was adequately represented. This study and others on cultivated flax diversity have revealed a significant genetic structure between fiber and oilseed groups. Further sub-structuring has also been characterized, often related to geographical origins or physiological development (winter vs. spring types) (Hoque et al., 2020; Fu et al., 2005; Speck et al., 2022). However, the effect of geographic origin is not always significant (Smýkal et al., 2011; Chandrawati et al., 2017; You et al., 2017). This may be attributed to the extensive exchange of genetic material (Soto-Cerda et al., 2013). Developing a core collection of flax germplasm focused on Western European diversity should facilitate genetic studies for flax breeding in Europe, while also allowing comparisons between studies based on this core collection.

In this study, we (i) performed genetic diversity analyses of a flax collection and (ii) compared various approaches to identify a core collection for further quantitative genetic studies. More specifically, we tested and evaluated approaches specifically designed for core collection construction alongside population optimization methods that were originally developed for genomic selection calibration sets. These methods differ in the type of input data used, the nature and number of optimization criteria (diversity indices, representativeness criteria, combination of them), and the algorithms used. The core collection designed will be useful for genome-wide association studies and genomic selection to enhance Western European flax breeding programs.

## 2 Materials and methods

### 2.1 Plant materials

The germplasm maintained by Arvalis since 2010 is a collection of 1,650 cultivated flax (fiber, oilseed and dual purposes type) accessions. The initial accessions were collected in 1938 from botanical collections and further extended through exchanges with research institutes such as INRAe, international biological resources centers, and breeding companies. The most recent accessions collected are lines originating from breeding programs and obtained in 2021. This diversity panel is predominantly composed of spring-type inbred lines, with 66% belonging to the oilseed group, 22% to the fiber group, and 12% classified as dual-purpose (both fiber and oilseed). Some winter-type lines have been included (oil and fiber) representing a valuable genetic source for low temperature tolerance. This germplasm encompasses the global diversity of cultivated flax, with accessions originating from 42 countries across all continents. The full list of accessions can be found in Supplementary Table S1.

### 2.2 Phenotypic data

The germplasm has been phenotyped for a set of 23 traits, summarized in Supplementary Table S2. These data are primarily passport data used to describe the accessions, including flower morphology (anther and pollen color, petal shape and color, filament color, style color, ciliation and coloration of capsule septa, corolla shape and size), seed morphology (seed color, thousand kernel weight), geographic origin, cultivated group (oilseed *versus* fiber-type), tolerance to low temperatures, lodging tolerance, as well as resistance to powdery mildew and Fusarium wilt. Prior to analysis, missing values were imputed using the R package missForest v1.5 (Stekhoven and Bühlmann, 2012). No imputation was performed for the country of origin.

### 2.3 Genotypic data

A seedling was produced for each of the 1650 flax accessions in the growing room at Arvalis Institute site in Boigneville (France) with 20°C/18°C day/night temperature. The Fresh leaves of two-weeks-old seedlings (50 - 100 mg) were harvested in microtube strips and flash-frozen at −80°C for 24 hours before being freeze-dried for 48 hours and then ground using the MM400 vibro-grinder (Retch). Genomic DNA was extracted from the crushed material using a modified Machery-Nagel NucleoMag Plant kit on the Beckman Coulter Biomek i5 automated workstation. Genomic DNA was then checked for quality on NanoDrop ND8000 (Thermo Fisher Scientific) and quantified on Qubit (Thermo Fisher Scientific) by Picogreen dosage. All accessions were genotyped using the Allegro AT-SNP-30K targeted genotyping tool (Demenou et al., 2025; Demenou et al., in preparation) at the EPGV platform (INRAe, Evry, France).

The genotyping matrix was generated through bioinformatic analysis, and then filtered. Markers and accessions with more than 50% missing data were discarded. The remaining markers were imputed using Beagle v5.4 (Browning, 2008; Browning and Browning, 2016), applying default parameters. Following this imputation, the genotyping matrix was filtered to remove markers having low minor allele frequency (MAF), retaining only those with MAF > 1%. This threshold has been chosen to preserve rare alleles that may carry valuable genetic information (Goudet et al., 2018). The distribution of selected imputed markers across the fifteen flax chromosomes was visualized using the R package CMplot v4.5.1 (Yin et al. 2021) to assess the quality and uniformity of the genotyping data.

### 2.4 Population structure and diversity analysis

Prior to the genetic diversity analysis, the genotyping matrix was intentionally pruned to retain only independent markers, thereby minimizing the confounding effects of collinearity among linked loci (Patterson et al., 2006). Marker pruning was performed using PLINK v1.07 (Purcell et al., 2007) with the following parameters: the ‘indep-pairwise’ function, a sliding window of 50 SNPs, and a linkage disequilibrium threshold of R² = 0.4.

We performed a Discriminant Analysis of Principal Components (DAPC) using the R package adegenet v2.1.10 (Jombart et al., 2010). DAPC assigns membership probabilities to predefined genetic clusters, which were inferred via K-means clustering. The number of retained principal components for the DAPC was determined using the Tracy-Widom test (Patterson et al., 2006), which identifies the first axes that significantly explain genetic variation. The optimal number of clusters K was determined by evaluating models with K-values ranging from 1 to 10, using the Bayesian Information Criterion (BIC) to select the best-supported model. Additionally, the most likely K-value was inferred by considering the correspondence between the identified groups and our germplasm knowledge. A Principal Component Analysis (PCA) was conducted on the pruned and standardized matrix using the R package FactoMineR v2.11 (Lê et al., 2008) to visualize the diversity and clustering. Pairwise Fixation indices (Fst) were calculated between genetic clusters using the R package hierfstat v0.5.11 (Goudet, 2005).

To further characterize the diversity hosted by the germplasm, expected mean heterozygosity (Berg and Hamrick, 1997), Shannon’s diversity index (Shannon, 1948), average Rogers’ genetic distance (Rogers, 1972), and the proportion of rare alleles (considering MAF < 0.10) were computed for each cluster and for all the entire germplasm.

### 2.5 Establishment of core-collections

Two categories of methods have been employed in this study: those specifically dedicated to core collections, and those aimed at building calibration populations, particularly for genomic selection purposes. Core collections were established using the R packages CoreCollection v0.9.5 (Jansen and Van Hintum, 2007; Odong et al., 2013), corehunter III v3.2.3 (Thachuk et al., 2009; De Beukelaer et al., 2018), TrainSel v3.0 (Akdemir et al., 2021), as well as the approach originally proposed by Laloë (1993) and further developed by Rincent et al. (2012) (R code acquired directly from the authors).

The method developed by Jansen and Van Hintum (2007) and later refined by Odong et al. (2013) is based on genetic distances among accessions. Entries are selected using a random descent algorithm, optimizing one of three available criteria: the Average Nearest Entry (A-NE), which minimizes the average distance between each accession and its nearest neighbors, the Nearest Neighboring Entry (E-NE), which maximizes this average distance, and the Entry-Entry (E-EE) criterion, which maximizes the pairwise distance among all accessions in the collection. We optimized the A-NE and E-NE criteria using Rogers’ genetic distance (Rogers, 1972). Optimization parameters were kept at their default settings.

The corehunter III R package (Thachuk et al., 2009; De Beukelaer et al., 2018) applies a stochastic local search algorithm based on replica exchange Monte Carlo chains for core collection development. Multiple selection criteria can be combined and weighted. This method can accommodate various input data types, including genetic distance matrices, genotypic and phenotypic datasets. Version III of this package supports the use of the following selection criteria, either individually or jointly: the previously described A-NE, E-NE, and E-EE criteria, expected heterozygosity (He), Shannon diversity index (SH), and allelic coverage (CV). All available optimization criteria were considered, except for the E-EE criterion. Criteria were applied individually or in pairwise combinations. In the case of bi-objective optimization, equal weights of 0.5 were assigned to each criterion. Additionally, a combination of the following three criteria, A-NE, SH, and CV, was tested, with each criterion assigned an equal weight (∼0.33). The execution mode was set to default, and normalization was applied for multi-objective optimizations.

The approach proposed by Laloë (1993) and elaborated by Rincent et al. (2012) aims to select a reference set of individuals for phenotyping that maximizes the reliability of genomic predictions for non-phenotyped individuals based on their genotypes. This method optimizes the generalized coefficient of determination (CDmean), which measures the correlation between predicted and observed values of genetic contrasts. CDmean balances the prediction error variance (PEV) against the genetic variance of the contrasts, accounting for genetic relatedness. The optimization is performed using a hill-climbing algorithm, exchanging one individual at each iteration, with the CDmean recalculated at every step using the individuals’ variance-covariance matrix. We use the R-code given by the authors. A total of 3,000 iterations were performed for each of the 10 core-collection replicates.

Other available tools for selecting calibration sets include STPGA (Akdemir et al., 2017), TSDFGS (Ou & Liao, 2019), and more recently, the R package TrainSel v3.0 (Akdemir et al., 2021). TrainSel enables the selection of individuals through mono- or multi-objective optimization, with possible weighting of criteria. It combines a genetic algorithm with simulated annealing. For our study, TrainSel was used with the following objective functions:

- D-optimality criterion (D-opt): aiming to maximize the determinant of the information matrix f(M), corresponding to the principal component transformation of the genotypic matrix, this criterion maximizes the dispersion in the multivariate genetic space;
- Avg_GRM_self: aiming to minimize the average relatedness within the calibration population, thus maximizing its genetic variance. The effectiveness of this criterion for calibration population selection has been demonstrated in previous studies (Atanda et al., 2021; Fernández-González et al., 2023);
- The combined optimization of D-opt and Avg_GRM_self;

Optimization algorithm hyperparameters were set as follows: medium population size, low complexity, and unordered sample. The remaining parameters were left at their default settings.

Table 1 summarizes the method × criterion combinations tested. For each combination, ten populations of 350 individuals were generated, with a random selection of the initial set. With such population size, the detection power for QTL studies should be enhanced (Hyne and Kearsey, 1995; Charmet, 2000; Vales et al., 2005).

**TABLE 1.**
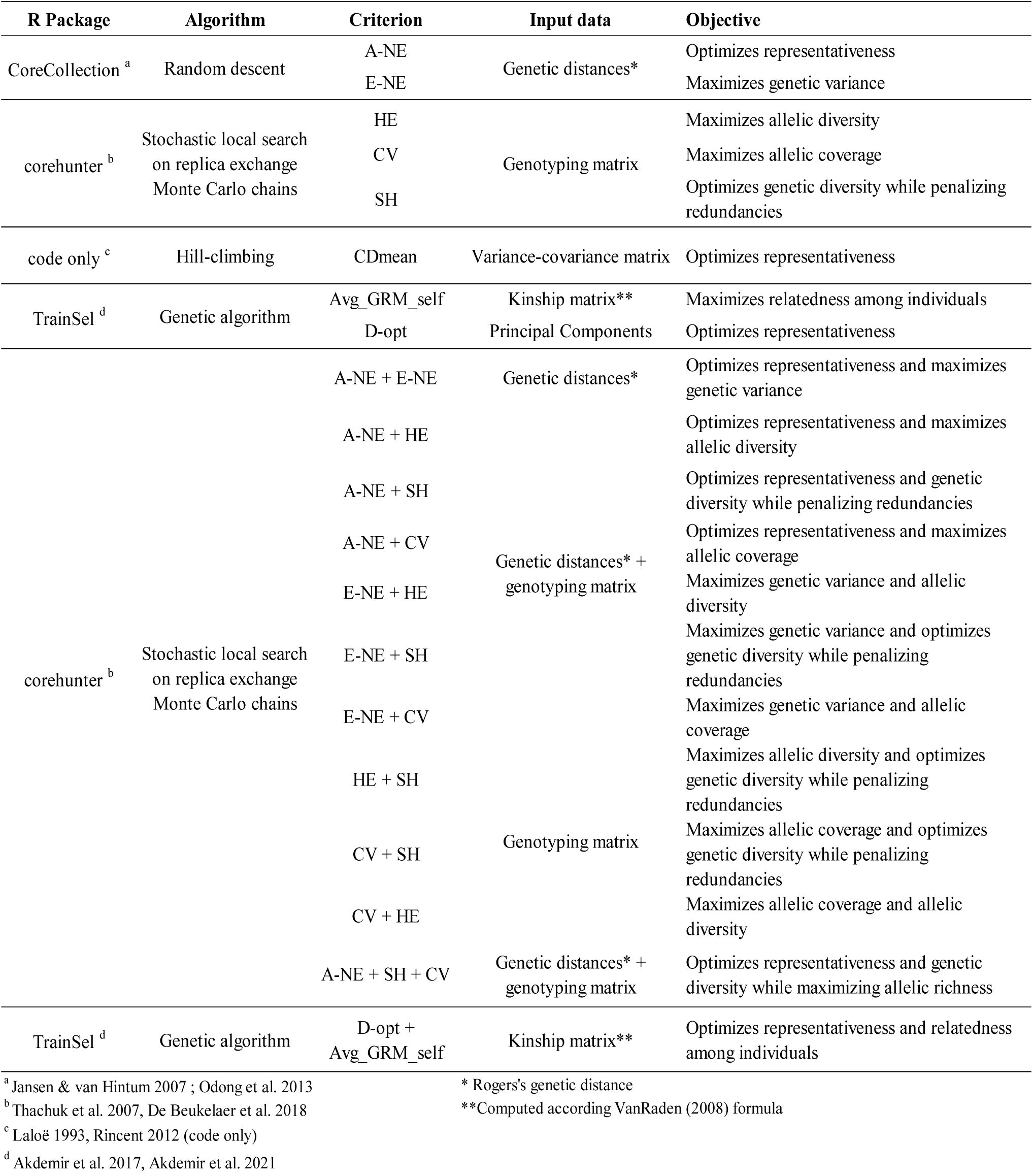
Combinations of twenty methods and selection criteria used in core collection development.

### 2.6 Core-collection evaluation

#### 2.6.1 Diversity and representativeness criteria

Each combination has been evaluated based on criteria assessing both genetic diversity and representativeness. To quantify the genetic diversity captured by each CC, the following metrics have been computed: the rare alleles ratio (MAF < 10%) (RAR), the mean Rogers’ distance (MRD), the expected heterozygosity (He), and the Shannon diversity index (SH). These indices are calculated using the following formulas:

1. Rare alleles ratio (RAR):

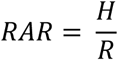 where R is the number of rare (i.e. MAF<10%) SNPs identified in the whole collection, and H the number of these rare SNPs founded as heterozygous within the core-collection.
2. Mean Rogers’ genetic distance (MRD) (Rogers, 1972):

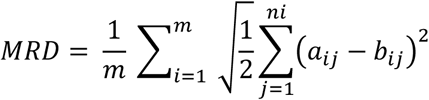 where m is the number of loci, n_i_ is the number of alleles at locus i, a_ij_ and b_ij_, are the genotype codes for individuals a and b at locus i. This metric can be likened to a Euclidean distance.
3. Expected heterozygosity (He) (Berg & Hamrick, 1997):

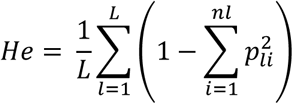 Where L is the number of loci, n_l_ is the number of alleles at locus l, p_li_ is the relative frequency of the i-th allele at locus l.
4. Shannon diversity index (SH) (Shannon, 1948):

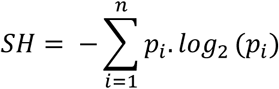 where n is the number of alleles and p_i_ is the frequency of the i-th allele.

To assess the representativeness of each CC relative to the WC, we computed the following metrics: allelic coverage (CV) (Kim et al., 2007), Kullback-Leibler divergence (KL) between allele frequency distributions of CC and WC (Kullback & Leibler, 1951), the average absolute Pearson’s correlation of principal component vectors (COR) between CC and WC (Yamamoto et al., 2007), and the Mean Difference ratio (MD) for a set of phenotypic variables (Hu et al., 2000). For MD calculation, independent phenotypic variables were preselected using Cramér’s V index (Cramér, 1999) to avoid overrepresentation of specific variable categories.

These representativeness metrics are computed using the following formulas:

1. Allelic coverage (CV):

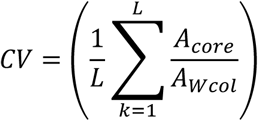 where L is the number of loci, A_core_ is the number of alleles present in the core collection at locus L and A_Wcol_ is the number of alleles present in the whole collection at the same locus.
2. Kullback-Leibler divergence (KL):

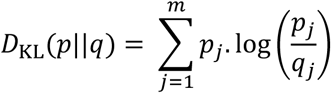 where m is the total number of SNPs, p_j_ is the frequency of the minor allele at SNP j in the core collection and q_j_ is the corresponding frequency in the whole collection.
3. Mean difference ratio for phenotypic traits (MD):

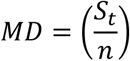 where S_t_ is the number of traits showing a significant difference between the CC and WC and n the total number of traits.
4. Average correlation between Principal Components (COR): Pearson correlation coefficients r_i_ are computed between principal components of the same rank from the CC and the WC. Due to an asymmetric distribution of correlation coefficients, we apply Fisher’s z-transformation (Fisher, 1921) before calculating the mean as follow:

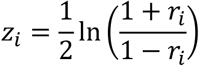 The value z_i_ is then transformed to obtain the average correlation:

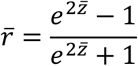 Since not all principal components contribute equally to the genetic variance, each transformed coefficient z_i_ is weighted by the eigenvalue (inertia) of the corresponding component in the WC. This weighting scheme assigns greater importance to the principal axes. Only the significant axes under the Tracy-Widom test (Patterson et al., 2006) are considered.

#### 2.6.2 QTL simulation and detection

We aimed to compare the core-collections on their ability to detect QTLs. To this end, we simulated two traits for each chromosome using the R package PhenotypeSimulator v0.3.4 (Meyer and Birney, 2018), using the genotypic matrix of the whole collection. The obtained simulated QTLs thus leverage the existing linkage disequilibrium. QTLs were simulated separately on each chromosome. To obtain QTLs evenly distributed along the genome, QTLs were thus simulated separately on each chromosome. In total, 940 QTLs distributed across the 15 flax chromosomes were obtained for the whole collection.

QTL detection was carried out using a mixed linear model (MLM) accounting for both population structure and relatedness (Yu et al., 2006). The model used was the following:

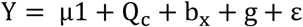

where Y is the vector of phenotypic simulated values, μ the overall mean, Q the matrix of covariates derived from the DAPC to capture population structure, c the vector of fixed effects associated with these covariates, b the additive fixed effect of the SNP, x the vector of SNP genotypes coded as 0, 1, or 2, g the vector of polygenic random effects, and ε the vector of residuals. Residuals were assumed to follow a normal distribution ∼ 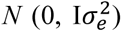, and polygenic effects were assumed to follow a normal distribution ∼ 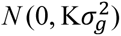, with K being the kinship matrix computed using the VanRaden (2008) method, as implemented in the R package AGHmatrix v2.1.4 (Amadeu et al., 2023). Mixed linear models were fitted using the R package GMMAT v1.4.2 (Chen et al., 2020). To assess the effectiveness of population structure correction, the genomic control inflation factor *λ* (Devlin and Roeder, 1999) was calculated for each trait. Values of *λ* below 1.05 were considered indicative of appropriate control for population structure effects (Price et al., 2010). SNP–trait associations with p-values below the significance threshold determined using Gao’s method (2008) were considered statistically significant and interpreted as putative QTLs. The proportion of QTLs identified within each core collection that were previously detected in the whole collection (considered as common QTLs) was calculated.

#### 2.6.3 Synthetic index for an appropriate comparison

To facilitate comparison among core-collection construction methods, we computed an index from the standardized values of our evaluation criteria. Criteria were pre-selected to balance representativeness and diversity. A preliminary analysis of the correlations between the indices was conducted to avoid including those that were highly redundant. The index *I* was defined accordingly as:

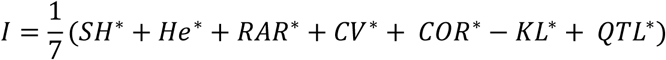

where SH is the Shannon index, He is the expected heterozygosity, RAR is the rare allele rate, CV is the allelic coverage, COR is the mean correlation coefficient, -KL is the negative Kullback-Leibler divergence and QTL is the proportion of shared QTLs. Each criterion has been standardized via normalization (i.e. centered and scaled).

## 3 Results

### 3.1 Genetic Diversity of the Flax Germplasm

A total of 30,893 Single Nucleotide Polymorphism (SNP) markers were obtained after genotyping the whole collection. Following filtering for missing data, the dataset comprised 28,475 SNP markers for 1,593 accessions with a minor allele frequency (MAF) greater than 1%. These markers are evenly distributed along the chromosomes, providing a significant genome-wide coverage (Figure 1). The imputed genotyping matrix was then pruned to retain only a set of independent markers more adapted for structure analyses. The resulting matrix contained 17,368 markers The distribution of Tracy-Widom test statistics (Supplementary Figure S3) indicated that the first 203 principal components significantly contributed to the genetic variance. These components were retained for the DAPC and subsequent analyses.

**FIGURE 1.**
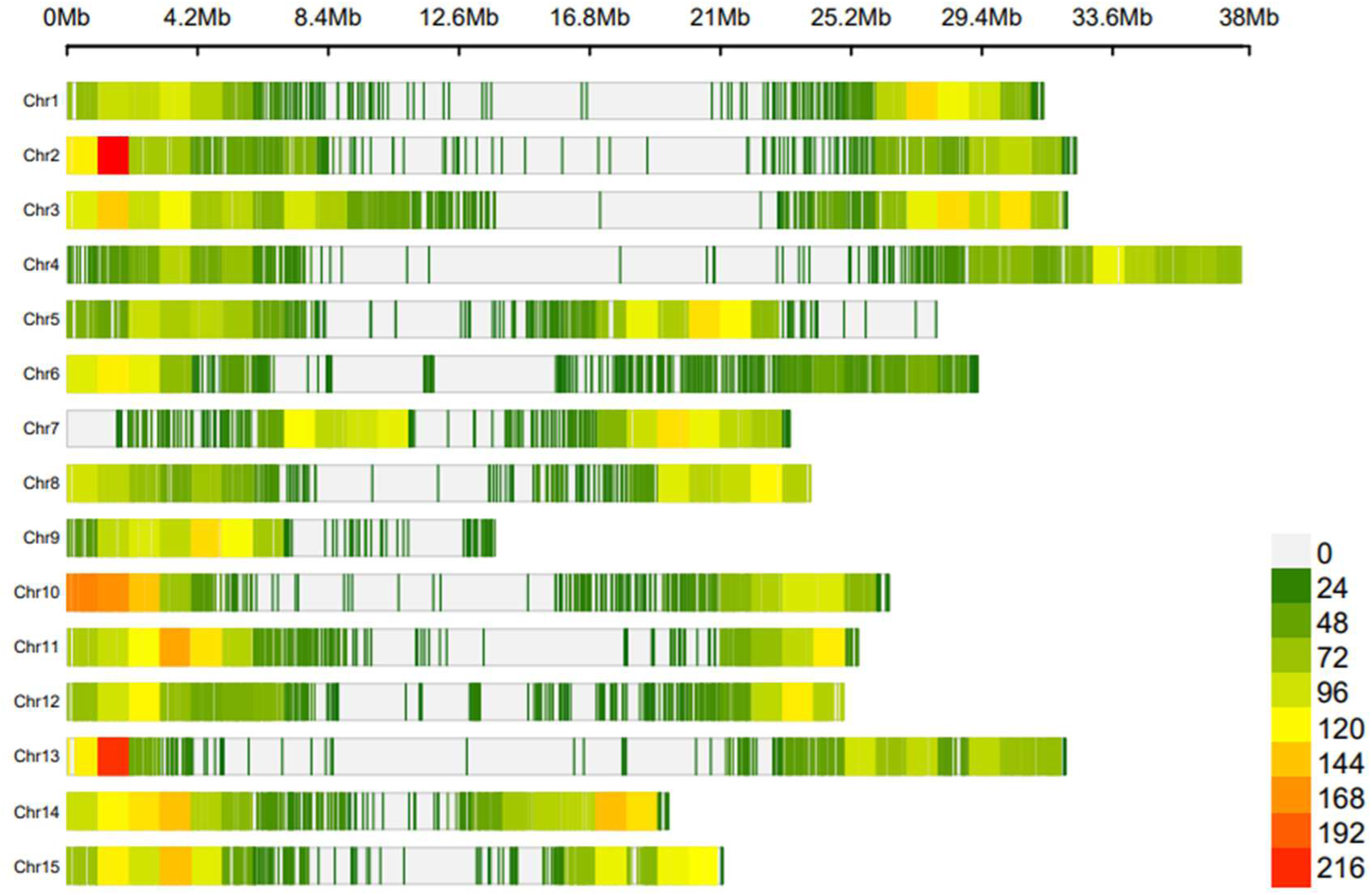
Distribution of SNP density along the 15 chromosomes of the flax germplasm, based on 28,475 mapped SNPs.

Based on the Bayesian Information Criterion (BIC), the most likely number of genetic clusters was determined to be 6 (Figure 2A). At K = 2, the structuring of the germplasm followed the cultivation type, distinguishing oilseed from fiber flax accessions. At K = 3, a winter flax cluster emerged, characterized by enhanced tolerance to low temperatures (data not shown). From K = 4 onwards, the structuring primarily reflected the geographical origin of accessions. For instance, at K = 4, a new cluster was identified within the oilseed group, separating South American flax accessions from the rest. At K = 5, another cluster was found within the fiber group, separating Western European fiber flax from Eastern European fiber flax. At K = 6, the oilseed group further subdivided into three sub-clusters: South American, North American, and Eastern European oilseed flax. Beyond K = 6, further differentiation occurred within the oilseed group, notably separating Eastern European lines from those of South American origin. Genetic diversity for K = 6 is illustrated via a principal component analysis (Figure 2B).

**FIGURE 2.**
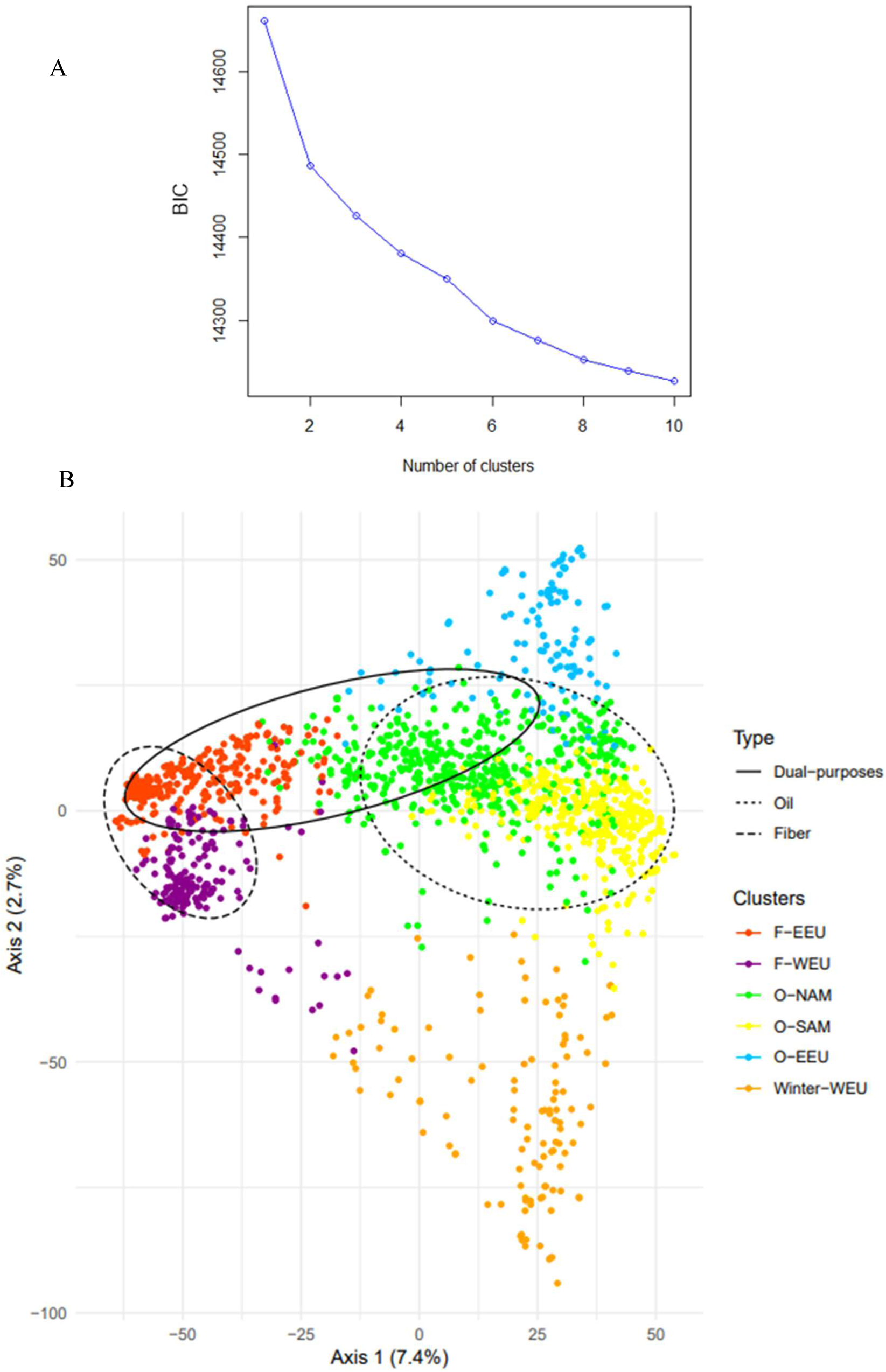
(A): Bayesian Information Criterion (BIC) values as a function of the number of clusters defined in the flax germplasm; (B) Scatter plot of the first two principal components of the PCA of the Arvalis flax germplasm (1,593 accessions), with color-coded clusters identified by DAPC.

Table 2 summarizes the main features of the six previously identified clusters. Oilseed lines are overrepresented relative to fiber lines (69% vs 31%). Cluster O-NAM contained the largest number of accessions (31%), mostly composed of North American spring-type oilseed flax. Cluster Winter-WEU contained the fewest accessions (7%), primarily consisting of winter flax originated from Western Europe.

**Table 2:** Genetic diversity and differentiation indices of the six identified flax clusters.

| Cluster | N | Ratio (%) | Diversity indices |  |  |  | Fixation index |  |  |  |  |
| --- | --- | --- | --- | --- | --- | --- | --- | --- | --- | --- | --- |
| | | | Shannon Index (SH) | Expected Heterozygosity ($H_e$) | Mean Rogers Distances (MRD) | Rare Alleles Ratio (RAR) | O-SAM | O-NAM | Winter WEU | O-EEU | F-EEU |
| O-SAM | 357 | 22% | 0.64 | 0.26 | 0.40 | 0.95 | - | - | - | - | - |
| O-NAM | 498 | 31% | 0.68 | 0.27 | 0.43 | 0.99 | 0.05 | - | - | - | - |
| Winter-WEU | 117 | 7% | 0.55 | 0.21 | 0.34 | 0.91 | 0.16 | 0.14 | - | - | - |
| O-EEU | 121 | 8% | 0.61 | 0.24 | 0.39 | 0.85 | 0.14 | 0.08 | 0.22 | - | - |
| F-EEU | 321 | 20% | 0.45 | 0.17 | 0.28 | 0.80 | 0.24 | 0.14 | 0.28 | 0.24 | - |
| F-WEU | 179 | 11% | 0.43 | 0.15 | 0.27 | 0.73 | 0.25 | 0.17 | 0.27 | 0.26 | 0.13 |
| Whole Collection | 1593 | 100% | 0.67 | 0.22 | 0.42 | 1.00 | 0.19 |  |  |  |  |

The pairwise fixation index Fst values revealed relatively high and significant genetic differentiation between clusters, especially between oilseed and fiber clusters (Fst up to 0.28, Table 2). The highest pairwise Fst value (Fst = 0.28) was found between clusters Winter-WEU and F-EEU (Western European oilseed vs. Eastern European fiber flax). The lowest Fst value (Fst = 0.05) was observed between oilseed clusters O-SAM and O-NAM, both comprising American oilseed lines. The Fst value between all clusters was significantly larger than zero (Fst =0.19).

Genetic diversity indices were generally moderate, with the oilseed clusters being the most diverse. (Table 2). The clusters exhibiting the greatest genetic diversity were the American oilseed lines (clusters O-SAM and O-NAM), with Shannon diversity indices ranging from 0.64 to 0.68 and expected heterozygosity ranging from 0.26 to 0.27. These clusters also harbored the highest rate of rare alleles. Conversely, the fiber clusters (F-EEU and F-WEU) exhibited the lowest level of diversity (He <0.17 and SH <0.45).

### 3.2 Core-collection comparison

We evaluated 20 *methods x selection* criteria combinations for core collections establishment. In total, 200 core collections, each consisting of 350 accessions, were generated and assessed. For each core collection, representativeness and diversity indices were computed. Additionally, the proportion of shared QTLs with the whole collection was measured. Table 3 summarizes the mean values (calculated from ten replicates) and standard deviation of the evaluation criteria for the twenty tested combinations (non-normalized data)..

**TABLE 3.** Evaluation criteria and standard errors computed for all core collection construction methods. SH: Shannon diversity index; He: Expected heterozygosity; MRD: Mean Rogers’s distances; RAR: Rare allele ratio (considering MAF<10%); CV: Allelic coverage; KL: Kullback-Leibler divergence of allelic frequencies; COR: Average correlation between Principal components; MD: Mean phenotypic differences; QTL: Ratio of common QTLs simulated.

| Methods | Diversity |  |  |  |  |  |  |  | Representativeness |  |  |  |  |  |  |  | QTL |  |
| --- | --- | --- | --- | --- | --- | --- | --- | --- | --- | --- | --- | --- | --- | --- | --- | --- | --- | --- |
|  | SH | sd (SH) | He | sd (He) | MRD | sd (MRD) | RAR | sd (RAR) | CV | sd (CV) | KL | sd (KL) | COR | sd (COR) | MD | sd (MD) | QTL | sd (QTL) |
| CDmean | 0.68 | 1.E-03 | 0.21 | 6.E-04 | 0.43 | 9.E-04 | 1.000 | 0.E+00 | 0.981 | 2.E-03 | 3.E-04 | 3.E-05 | 0.65 | 2.E-02 | 0.11 | 0.06 | 27% | 0.04 |
| CV | 0.67 | 3.E-03 | 0.21 | 3.E-03 | 0.42 | 2.E-03 | 1.000 | 0.E+00 | 0.992 | 1.E-03 | 4.E-04 | 4.E-05 | 0.59 | 2.E-02 | 0.12 | 0.06 | 27% | 0.02 |
| Avg_GRM | 0.67 | 3.E-03 | 0.21 | 2.E-03 | 0.43 | 2.E-03 | 1.000 | 2.E-04 | 0.991 | 1.E-03 | 4.E-03 | 1.E-02 | 0.58 | 4.E-02 | 0.17 | 0.05 | 27% | 0.03 |
| ANE_CV | 0.67 | 2.E-03 | 0.21 | 3.E-03 | 0.42 | 2.E-03 | 1.000 | 8.E-05 | 0.992 | 1.E-03 | 4.E-04 | 5.E-05 | 0.57 | 4.E-02 | 0.14 | 0.07 | 28% | 0.03 |
| ANE | 0.68 | 3.E-03 | 0.21 | 2.E-03 | 0.43 | 2.E-03 | 1.000 | 5.E-05 | 0.991 | 1.E-03 | 6.E-04 | 2.E-04 | 0.56 | 3.E-02 | 0.13 | 0.05 | 26% | 0.04 |
| D-opt+Avg_GRM | 0.68 | 2.E-03 | 0.21 | 1.E-03 | 0.43 | 2.E-03 | 1.000 | 4.E-04 | 0.985 | 2.E-03 | 2.E-02 | 4.E-02 | 0.51 | 3.E-02 | 0.16 | 0.06 | 26% | 0.02 |
| D-opt | 0.69 | 1.E-03 | 0.21 | 1.E-03 | 0.44 | 8.E-04 | 1.000 | 1.E-04 | 0.965 | 2.E-03 | 1.E-03 | 1.E-04 | 0.50 | 1.E-02 | 0.22 | 0.08 | 28% | 0.02 |
| ENE | 0.69 | 1.E-03 | 0.22 | 4.E-03 | 0.44 | 1.E-03 | 1.000 | 2.E-04 | 0.994 | 1.E-03 | 2.E-03 | 3.E-04 | 0.44 | 4.E-02 | 0.34 | 0.12 | 24% | 0.02 |
| ANE_ENE | 0.69 | 4.E-04 | 0.21 | 7.E-04 | 0.44 | 3.E-04 | 0.999 | 2.E-04 | 0.974 | 1.E-03 | 3.E-03 | 1.E-04 | 0.41 | 7.E-03 | 0.49 | 0.08 | 18% | 0.01 |
| SH_CV | 0.71 | 1.E-04 | 0.22 | 9.E-04 | 0.46 | 1.E-04 | 1.000 | 0.E+00 | 0.991 | 5.E-04 | 6.E-03 | 6.E-05 | 0.41 | 9.E-03 | 0.49 | 0.04 | 27% | 0.02 |
| ANE_SH | 0.71 | 1.E-04 | 0.22 | 1.E-03 | 0.45 | 1.E-04 | 0.999 | 4.E-04 | 0.991 | 4.E-04 | 2.E-02 | 4.E-02 | 0.48 | 6.E-03 | 0.43 | 0.04 | 24% | 0.04 |
| ANE_SH_CV | 0.71 | 4.E-04 | 0.22 | 9.E-04 | 0.45 | 3.E-04 | 1.000 | 0.E+00 | 0.991 | 7.E-04 | 4.E-03 | 1.E-04 | 0.48 | 2.E-03 | 0.39 | 0.04 | 20% | 0.01 |
| ANE_HE | 0.71 | 1.E-04 | 0.22 | 1.E-03 | 0.45 | 8.E-05 | 0.998 | 3.E-04 | 0.991 | 5.E-04 | 1.E-01 | 1.E-02 | 0.46 | 3.E-03 | 0.46 | 0.03 | 20% | 0.03 |
| SH | 0.71 | 2.E-05 | 0.22 | 7.E-04 | 0.46 | 1.E-04 | 0.997 | 4.E-04 | 0.990 | 5.E-04 | 1.E-01 | 3.E-03 | 0.41 | 6.E-03 | 0.51 | 0.06 | 27% | 0.02 |
| HE_SH | 0.71 | 5.E-05 | 0.22 | 9.E-04 | 0.46 | 5.E-05 | 0.996 | 5.E-04 | 0.990 | 2.E-04 | 1.E-01 | 5.E-03 | 0.39 | 1.E-02 | 0.51 | 0.04 | 27% | 0.01 |
| HE_CV | 0.71 | 7.E-05 | 0.22 | 2.E-03 | 0.46 | 1.E-04 | 1.000 | 0.E+00 | 0.991 | 8.E-04 | 7.E-03 | 8.E-05 | 0.38 | 8.E-03 | 0.54 | 0.05 | 26% | 0.01 |
| ENE_SH | 0.71 | 1.E-04 | 0.22 | 3.E-04 | 0.45 | 1.E-04 | 0.998 | 2.E-04 | 0.982 | 1.E-03 | 1.E-01 | 2.E-04 | 0.37 | 9.E-03 | 0.53 | 0.03 | 21% | 0.02 |
| HE | 0.71 | 7.E-05 | 0.22 | 1.E-03 | 0.46 | 1.E-04 | 0.995 | 7.E-04 | 0.990 | 5.E-04 | 1.E-01 | 4.E-03 | 0.35 | 3.E-02 | 0.51 | 0.05 | 25% | 0.02 |
| ENE_HE | 0.71 | 1.E-04 | 0.23 | 1.E-03 | 0.45 | 1.E-04 | 0.995 | 8.E-04 | 0.982 | 9.E-04 | 1.E-01 | 8.E-04 | 0.18 | 2.E-02 | 0.57 | 0.05 | 19% | 0.02 |
| ENE_CV | 0.69 | 3.E-04 | 0.22 | 2.E-03 | 0.44 | 2.E-04 | 1.000 | 0.E+00 | 0.975 | 1.E-03 | 5.E-03 | 2.E-04 | 0.32 | 9.E-03 | 0.49 | 0.07 | 21% | 0.02 |

The evaluation criteria showed different levels of variability between the methods. The Rare Allele Ratio (RAR) exhibited the lowest variability across methods. Regardless of the method used, nearly all rare alleles were consistently captured. Similarly, the allelic coverage rate (CV) displayed limited variation across methods (ranging from 0.965 to 0.994), suggesting that these two metrics were not strongly discriminative. In contrast, diversity-related indices such as Shannon’s index (SH), expected heterozygosity (He), and Mean Rogers ‘Distances (MRD) demonstrated more substantial variation and followed similar trends across methods (Table 3). As expected, methods that maximized these indices tended to yield higher overall genetic diversity. Representativeness indices such as Kullback-Leibler divergence (KL), correlation coefficient (COR), and Mean Differences (MD) revealed significant differences between methods. The CDmean method consistently achieved the lowest KL divergence, the highest COR, and the lowest MD values. Other methods showing high representativeness included the CV, Avg_GRM_self, and ANE-based approaches. In general, representativeness indices exhibited greater variability compared to diversity indices. The highest MD values (reflecting low phenotypic representativeness) were observed for methods such as ENE_HE, He_CV, and ENE_SH.

All QTL detection models successfully controlled the inflation of test statistics (see Supplementary figure S4a & S4b). QTL detection varied between methods, ranging from 18% to 28% of the 940 QTL detected in the whole collection (Supplementary figure S5). Methods that detected the highest average number of simulated QTLs included D-opt, ANE_CV, and SH_CV. CDmean also showed a high median catching rate, ranking second only to D-opt. The ANE_CV, ANE_He and ANE_SH methods were more subject to sampling variations, exhibiting greater variability in QTL detection rate. Generally, methods ensuring high representativeness tended to catch more simulated QTLs. The method with the lowest QTL rate is ANE_ENE, and methods that prioritized the ENE index tended to bring fewer QTLs overall.

A clear trade-off was observed between maximizing diversity and maximizing representativeness. Methods that were most effective at enhancing diversity generally performed less in terms of representativeness, and vice versa. Nonetheless, certain methods, such as ANE_SH, ANE_SH_CV, and SH_CV, provided a balanced compromise between the two objectives.

### 3.3 Composite score index for core collection evaluation

For a simplified cross-method comparison, a composite index integrating some evaluation criteria was calculated. Prior to index construction, inter-criteria correlations were assessed to avoid the inclusion of highly collinear metrics. Concurrently, a balance between representativeness and diversity was sought. The MD index was intentionally excluded from this composite index because it relies primarily on passport data that cannot accurately reflect the full extent of phenotypic diversity. Correlation analysis (Supplementary Figure S6) revealed that diversity criteria (SH, MRD, and He) were mutually and significantly correlated; moreover, MRD and SH exhibited redundancy, warranting the inclusion of only one of these metrics in the composite index. Although KL and COR were correlated, they conveyed distinct information and were therefore both retained. The distributions of KL, RAR, and CV were found to be highly skewed (Supplementary Figure S6). Notably, COR exhibited the strongest correlation with the QTL criterion (R = 0.45). All the seven selected criteria were normalized before integration into the composite score. The resulting composite index values are summarized in the boxplot Figure 3.

**FIGURE 3.**
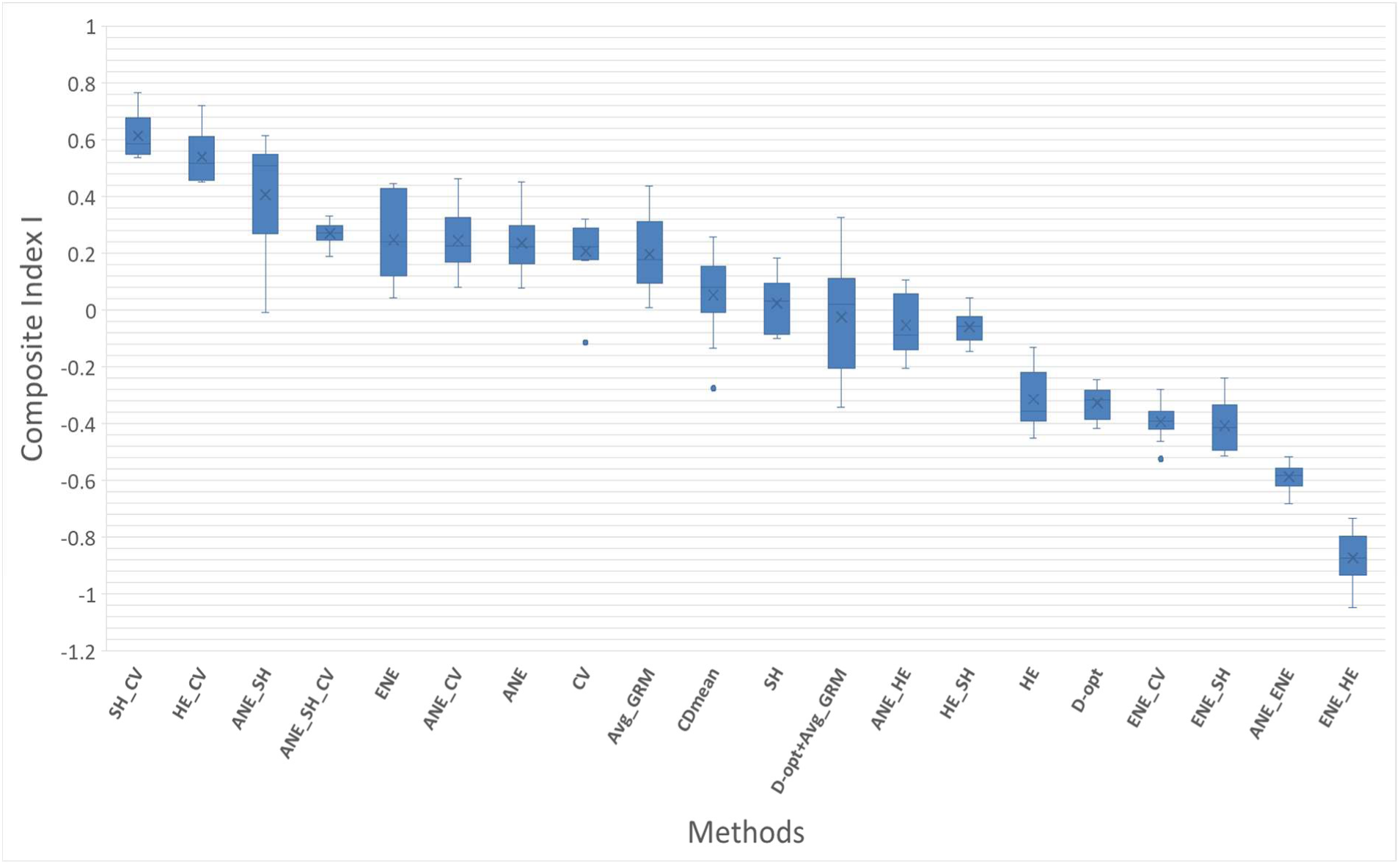
Boxplot of the composite score index values (I) calculated across all tested methods for core collection development. For each method, 10 core collection have been generated. The crosses (X) illustrate median values. The first and third quartiles and the outliers are represented.

The method yielding the highest index value, and thus the best overall compromise was the one that simultaneously optimized both the SH and CV criteria (SH_CV). In general, methods that integrated multiple criteria tended to produce higher index scores.

Conversely, the method with the lowest index was the one that combined ENE and HE optimization (ENE_HE). More broadly, methods that optimized the ENE criterion, either alone or in combination with other criteria, consistently resulted in lower index values.

## 4 Discussion

### Population Structure and Genetic Diversity

The genetic diversity study of the worldwide flax collection revealed a pronounced genetic structure within this germplasm. The primary axis of differentiation was between oilseed flax and fiber flax types as expected, a pattern consistent with previous findings reported in the literature (Fu et al., 2005; Hoque et al., 2020; Speck et al., 2022). At K=3, the emergence of a new cluster composed predominantly of winter-type flax lines highlights the differentiation between spring and winter types. This type of structuring has also been observed in previous studies (Uysal et al., 2010; Fu, 2012; Hoque et al., 2020). This cluster predominantly comprised lines tolerant to low temperatures and thus represents a valuable reservoir of alleles for the improvement of other clusters, particularly given its composition of both fiber and oilseed genotypes. For K=4, the population structure became more refined, with subdivisions reflecting the geographical origin of the accessions. A new cluster composed primarily by south American oilseed lines appeared. In contrast, the fiber flax group, only began to subdivide from K=5 onwards, distinguishing between Western and Eastern European origins. At K=6 and for higher K values, further subdivision occurred within the oilseed clusters, notably distinguishing Eastern European flax from Western European and South American oilseed flax. In the K-means clustering procedure, although the Bayesian Information Criterion exhibited a peak at K=2, it ultimately designated six clusters as the optimal configuration. The investigation of cluster structure could be supplemented by the Bayesian framework of Pritchard *et al*. (2000) using the STRUCTURE software, thereby reinforcing our results.

Genetic diversity analyses for the whole collection revealed moderate diversity (*H* e= 0.22), a value similar to that reported by Hoque et al., (2020) who used 6200 SNP markers to analyse the diversity of 350 genotypes. This level of diversity is expected, given that flax possesses an autogamous reproductive system (Hoque et al., 2020). The expected heterozygosity value of the clusters revealed that oilseed flax clusters harbored greater diversity than fiber flax clusters (Table 2). These findings are consistent with those reported in the literature. Hoque et al., 2020 reported seven genetic clusters with only one cluster for fiber type flax. This difference is probably due to the history of domestication and selective breeding focused on specific traits in each type. Oil flax is considered the ancestral form from which fiber flax was derived. During domestication, fiber flax underwent strong selection for traits like stem length and fiber quality, which reduced its genetic diversity compared to oil flax (Xie et al, 2018). Selective breeding for specific fiber traits in fiber flax led to narrowing the gene pool. Genomic studies confirmed that many genes associated with fiber traits in fiber flax showed strong selection signals, further supporting the idea of a genetic bottleneck (Povkhova et al., 2021). Furthermore, fiber flax is cultivated in a more restricted geographic area compared to oilseed flax, thus leading to high selective pressure to adapt varieties to the specific agro-climatic conditions of this growing area. In contrast, oil flax retained more of the original genetic variation because it was selected for a broader range of traits, including oil content and seed characteristics (Jiang et al., 2021). This finding further underscores that oilseed flax lines harbor a more substantial diversity reservoir, a factor that likely accounts for their predominance over fiber flax lines in conservation collections.

### Core-collection Assessment

In this study, we evaluated a comprehensive suite of core-collection development approaches (twenty method x selection-criterion combinations) resulting in 200 core collections of 350 accessions each. Dedicated core-collection methods were compared both among themselves and against calibration-population optimization approaches (as for genomic selection calibration methods). Our aim was to assess outcomes from approaches that differ in their input data (e.g., diversity indices, genetic-distance matrices, or kinship matrices) and in their optimization criteria. The underlying algorithms also varied between methods.

First, we observed that the evaluation criteria did not exhibit the same level of variability. Both the CV and RAR criteria showed low variability across methods. All approaches managed to capture most alleles, including the rarest. This limited variability may be attributed to the core collections size. Indeed, with 350 individuals selected, it is more likely to encompass the full allelic diversity of the initial collection of 1,593 individuals. It would be relevant to compare the tested methods using smaller core collections, ranging from 50 to 100 individuals for example. With such reduced sample sizes, differences between methods might become more pronounced. The choice of using 350 individuals was based on studies assessing the statistical power for QTL detection (Hyne and Kearsey, 1995; Charmet, 2000; Vales et al., 2005) but also for practical reasons with the perspective of testing this CC in field experiments.

Secondly, we observed that diversity-related criteria exhibited lower variability across methods compared to representativeness-related criteria. With core collections composed of 350 individuals, genetic diversity is rapidly captured. This sample size corresponds to approximately 22% of the total population, which exceeds the proportion of 10% generally recommended in the literature (Van Hintum, 2000). Nevertheless, methods that jointly optimized He with CV, or SH with CV, significantly increased genetic diversity within the core collections compared to other methods.

Among the methods that best preserved representativeness, CV, ANE, ANE_CV, Avg_GRM_self, and CDmean produced core collections that closely mirrored the initial collection. These methods effectively maintained the overall genetic structure. As expected, the methods originally developed for optimizing calibration populations (CDmean and Avg_GRM_self) yielded core-collections that were highly representative of the whole collection. Notably, high levels of representativeness can be achieved through different strategies, by maximizing allelic coverage, minimizing the average distance between an accession and its nearest neighbors (ANE approach), or by optimizing pairwise relatedness between individuals.

Regardless of the approach, a clear trade-off emerged between optimizing representativeness and maximizing the intrinsic diversity of the core collection. Optimizing for diversity can lead to over-representation of rare alleles, which may not reflect the typical characteristics of the full germplasm, thus reducing representativeness (Marita et al., 2000; Franco-Duran et al., 2019). Conversely, optimizing for representativeness may result in a CC that misses rare alleles or unique genotypes, thus underrepresenting the full spectrum of the diversity (De Beukelaer et al., 2018). Some advanced algorithms attempt to balance both objectives, but improving one often comes at the expense of the other, requiring a compromise based on the intended use of the core-collection.

In our study, we also sought to assess the capacity of core collections to capture QTLs. Core collections are particularly valuable for QTL discovery because they harbor extensive genetic diversity and thus represent a rich source of QTLs of interest (SotoCerda et al., 2014; McLeod et al., 2023). It is therefore important to determine which optimization criteria yield a core collection that maximizes QTL detection power. We detected on average 225 QTLs out of the 940 simulated on the whole collection, corresponding to approximately 24% overlap in detected QTLs. This reduction in detection, observed regardless of the method employed, can be attributed to the smaller size of the core collections, which diminishes power to detect QTLs with smaller effect sizes. Indeed, numerous studies have demonstrated that QTL detection power is strongly influenced by the population size: larger populations consistently achieve higher detection power revealing more QTLs, especially those with minor effects, whereas small populations often fail to detect these minor QTLs (Vales et al., 2005; Wang et al., 2012; Wang and Xu, 2019; Nwogwugwu et al., 2022). Among the methods tested, the highest number of QTLs detected within a core collection was obtained using the ANE_CV method, where one core-collection allowed detecting a total of 299 QTLs. However, this method exhibited high variability in QTL detection rates. Such variability arised because each random seed initiates the selection with a different individual, leading to a distinct ensemble of cluster centers. The ANE method’s combination of cluster center representativeness (ensuring thorough coverage of each region in genotype space) and randomized starting points (inducing different traversals of that space) naturally produces subpopulations with unique allele frequency landscapes and linkage patterns. Since QTL detection critically depends on these landscapes and patterns, ANE yields high variability in the sets of QTLs discovered across different core-collections. The D optimality criterion method (Dopt) create CC that capture on average the highest number of QTLs, with relatively low variability across replicates giving therefore a more stable QTL detection. The Dopt method selects a subset of individuals that optimally represents the genetic diversity and structure of the whole collection. By maximizing the determinant of the feature matrix (typically the first *q* principal components of the marker matrix), Dopt ensures that the selected subset spans the broadest possible range of genetic variation. This is crucial for QTL detection because a training set encompassing the full spectrum of genetic diversity increases the likelihood that QTL alleles segregate within the subset, thereby enhancing detection power. Moreover, maximizing the determinant reduces multicollinearity in the design matrix, yielding more precise estimates of marker effects. In general, methods that maximized representativeness, such as Dopt, ANE_CV, and Avg_GRM_self, captured the largest number of simulated QTLs. Any subpopulation that preserves the same underlying genetic structure of the whole collection necessarily has a higher probability of including those QTLs.

Evaluating methods based on multiple indices can make selecting the best approach challenging. To facilitate the identification of an optimal trade-off between diversity, representativeness, and QTL detection efficiency, we computed a composite index. This index integrated the criteria in a comprehensive and balanced manner. Significant differences in index values were observed across methods. In our study, the combined optimization of the Shannon index and the allelic coverage (with equal weighting) yielded the highest average composite index. The Shannon index favors both high allelic richness and evenness in allele frequencies, thereby promoting allelic balance. Balanced allele frequencies increase the likelihood that causal variants (QTLs) segregate at detectable frequencies. The allelic coverage criterion tends to generate populations that are structurally similar to the whole collection, thus enhancing representativeness. These two criteria appear to be complementary. We also observed that methods combining the ENE criterion with other criteria tended to yield lower average composite index values. The ENE criterion maximizes the genetic distances between the selected entry and its neighboring accessions. This tends to select individuals located at the extremes of the diversity cloud. Combinations including ENE did not produce good compromises in our evaluations.

Evaluating the quality of a core collection should, whenever possible, be based on data that were not used in its construction (van Hintum et al., 2000). In the present study, we employed the full set of SNP markers both to assemble the core collections and to assess their performance. A more impartial evaluation could be achieved by partitioning the marker dataset: one subset of independent, evenly spaced SNPs, would be used to define the core collections, and the remaining marker set would serve exclusively for their validation. This two-step approach would reduce circularity and provide a more rigorous assessment of core-collection construction methods.

## 5 Conclusion

The diversity analysis of the Arvalis flax germplasm revealed a moderate genetic diversity and a clear genetic split between oilseed and fiber types, with additional clusters reflecting seasonal and geographic variation. When reducing the germplasm to 350 accessions across twenty sampling strategies, most methods captured nearly all alleles but differed substantially in representativeness and QTL detection power. While ANE_CV detected the most QTLs, it showed high variability, and D-optimality offered a more stable and significant recovery. By integrating diversity, representativeness, and QTL-detection into a composite index, the Shannon-index plus allelic coverage (SH + CV) combination emerged as the superior compromise for our case study, maximally balancing genetic richness, representativeness, and trait-discovery potential for GWAS applications.

## Data availability statement

The genotyping methodology will be published (Demenou et al., in preparation). SNPs developed on the full germplasm can be found online.

## Author contributions

BD has elaborated the research project, performed the establishment of the flax germplasm and has provided genotypic and phenotypic data. MF has provided the purified DNA samples before genotyping. MG analyzed data and wrote the manuscript with significant input from all authors. All authors read and approved the final manuscript.

## Acknowledgments

The authors wish to thank, Isabelle CHAILLET (Arvalis Institute) and Christophe PINEAU (Linea Semences) for their help conceiving the CoreFLAX projet (Genetic diversity analysis of cultivated flax genetic resources and definition of flax core collection); and. They would also like to thank Adriane Rolland and Caroline Laffray from the GenoPAV laboratory (Arvalis) for producing the seedlings and extracting the DNA for the entire collection. Finally, they would like to thank Patricia Faivre-Rempant, Damien Hinsinger and the entire team at the INRAE Unit ‘Étude du Polymorphisme des Génomes Végétaux’ (EPGV) for their help with the genotyping of the entire collection.

