## Supplementary Figures S3 to S6 for "Optimizing core collections for genetic studies: a worldwide flax germplasm case study"

**Supplementary Figure S3**

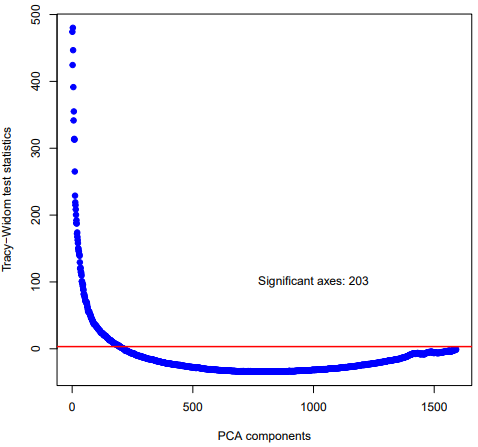

Tracy-Widom test statistics according to the principal components

**Supplementary Figure S4**

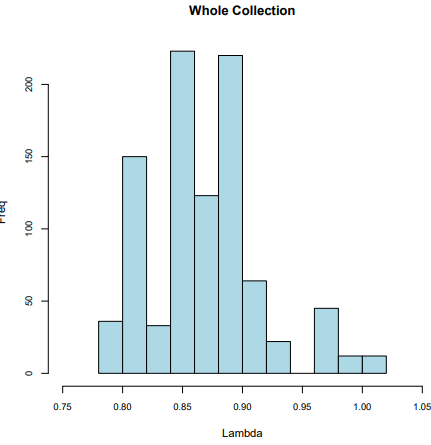

a) Genomic control inflation factor distribution computed for the whole collection after GWAS analysis

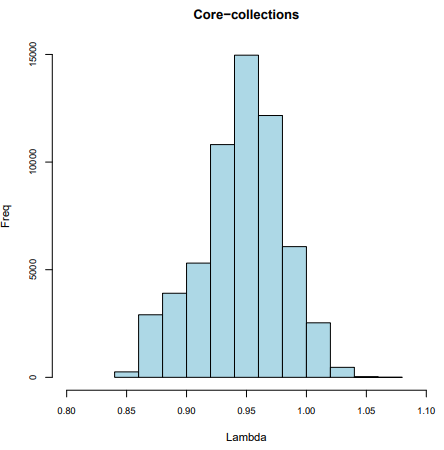

b) Genomic control inflation factor distribution computed for the 200 core-collections collection after GWAS analysis.

**Supplementary Figure S5**

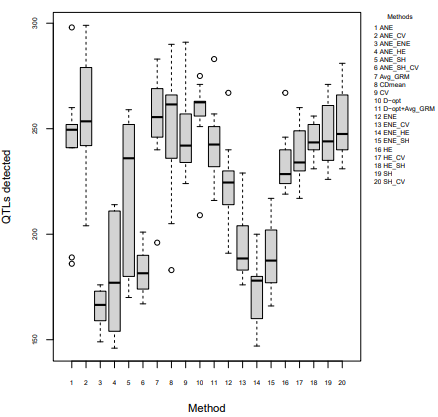

Number of detected QTLs based on core-collection creation methods. For each method tested, 10 core-collections were created. QTLs were simulated from the whole collection.

**Supplementary Figure S6**

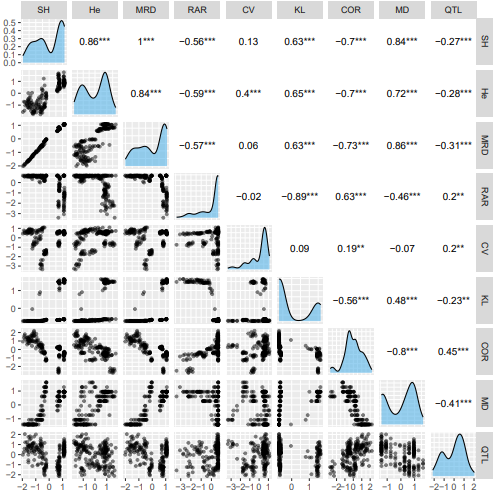

Distribution of the diversity and representativeness criteria and their pairwise Pearson’s correlation coefficient computed for the 200 core-collections created.
